# A High Avidity Biosensor Reveals PI(3,4)P_2_ is Predominantly a Class I PI3K Signaling Product

**DOI:** 10.1101/410811

**Authors:** Brady D. Goulden, Jonathan Pacheco, Allyson Dull, James P. Zewe, Alexander Deiters, Gerald R. V. Hammond

**Affiliations:** Department of Cell Biology, University of Pittsburgh School of Medicine and Pittsburgh, PA, USA; Department of Chemistry, University of Pittsburgh, Pittsburgh, PA, USA

**Keywords:** PtdIns(3,4)*P*_2_, inositol lipid, phosphoinositides

## Abstract

Class I PI 3-kinase (PI3K) signaling is central to animal growth and metabolism, and disruption of this pathway occurs frequently in cancer and diabetes. However, the specific spatial/temporal dynamics and signaling roles of its minor lipid messenger, phosphatidylinositol (3,4)-bisphosphate [PI(3,4)P_2_], are not well understood. This owes principally to a lack of tools to study this scarce lipid. Here, we developed a high sensitivity genetically encoded biosensor for PI(3,4)P_2_, demonstrating high selectivity and specificity of the sensor for the lipid. We show that despite clear evidence for class II PI3K in PI(3,4)P_2_-driven function, the overwhelming majority of the lipid accumulates through degradation of class I PI3K-produced PIP_3_. However, we show that PI(3,4)P_2_ is also subject to hydrolysis by the tumor suppressor lipid phosphatase PTEN. Collectively, our results show that PI(3,4)P_2_ is potentially an important driver of class I PI3K-driven signaling, and provides powerful new tools to begin to resolve the biological functions of this lipid downstream of class I and II PI3K.

## Introduction

Class I phosphoinositide 3-OH kinase (PI3K) signaling is central to the control of growth and metabolism in animals (Vanhaesebroeck et al., 2012). Overactivation of this pathway is the most common event in cancer (Fruman et al., 2017), yet given its major role in insulin signaling, inhibition of the pathway triggers insulin resistance and type 2 diabetes (Hopkins et al., 2018). Therefore, the ability to selectively manipulate PI3K signaling could have tremendous therapeutic benefit. Efforts to accomplish this goal are a major focus of the biomedical enterprise (Fruman et al., 2017)

At the molecular level, PI3K signaling involves the generation of the plasma membrane (PM) second messenger lipids phosphatidylinositol 3,4,5-trisphosphate (PIP_3_) and phosphatidylinositol 3,4-bisphosphate [PI(3,4)P_2_] that activate downstream effector proteins like the serine/threonine kinase Akt. PIP_3_ is the major lipid produced, and most functions of the pathway are attributable to it (Vanhaesebroeck et al., 2012). PI(3,4)P_2_ has instead been viewed as either a degradation product (Ishihara et al., 1999), or as an alternative activator of the pathway (Ebner et al., 2017). However, selective functions for PI(3,4)P_2_ have recently been described that are independent of PIP_3_ (Li and Marshall, 2015). These include the formation of lamellipodia and invadopodia (Krause et al., 2004; Bae et al., 2010; Oikawa et al., 2008; Sharma et al., 2013), along with clathrin-mediated and clathrin-independent endocytosis (Posor et al., 2013; Boucrot et al., 2015) In each case, these functions could conceivably be driven by, or occur independently of, class I PI3K signaling.

Synthesis of PI(3,4)P_2_ can proceed via three routes: In the first, class I PI3K directly generates PI(3,4)P_2_ and PIP_3_ by 3-OH phosphorylation of the respective PM phosphoinositides PI4P and PI(4,5)P_2_ (Carpenter et al., 1990). Subsequently, the observation that PI(3,4)P_2_ synthesis lags behind PIP_3_ in stimulated cells (Stephens et al., 1991; Jackson et al., 1992; Hawkins et al., 1992), coupled with the discovery of the PIP_3_-specific 5-phopshatase enzymes SHIP1 and SHIP2 (Damen et al., 1996; Pesesse et al., 1997), led to the proposal of a second route: PI(3,4)P_2_ production by removal of the 5-OH phosphate from PIP_3_. More recently, a third route has been established, again invoking direct phosphorylation of PI4P, this time by class II PI3K enzymes (Domin et al., 1997; Posor et al., 2013). However, the relative contributions of these pathways, and how they couple to disparate PI(3,4)P_2_-dependent cellular functions, remains unclear (Li and Marshall, 2015).

Resolving how the spatial/temporal dynamics of PI(3,4)P_2_ signaling couples to different biological functions requires approaches to identify the lipid in intact, living cells. Isolated lipid binding domains fused to fluorescent reporters often make highly selective genetically encoded biosensors for this purpose (Wills et al., 2018). The pleckstrin homology (PH) domain on the C-terminus of TAPP1 (TAndem Ph-domain containing Protein 1) exhibits specific binding to PI(3,4)P_2_ in the test tube (Thomas et al., 2001; Dowler et al., 2000). As a result, several studies have employed fluorescent protein conjugates of this domain to track PI(3,4)P_2_ signaling, though the domain fails to detect resting levels or the limited accumulation of the lipid in response to stimuli such as insulin-like growth factor (Kimber et al., 2002; Marshall et al., 2002; Oikawa et al., 2008; Posor et al., 2013).

Herein, we developed a higher-avidity tandem trimer of PH-TAPP1. We show PI(3,4)P_2_ generation is sufficient to recruit the probe, which is exquisitely selective for the lipid over other phosphoinositides. We then demonstrate that the class I PI3K pathway, acting via PIP_3_ synthesis, dominates PI(3,4)P_2_ accumulation in cells. Our data also support the recently proposed direct degradation of both PI(3,4)P_2_ and PIP_3_ by the lipid phosphatase and tumor suppressor PTEN (Malek et al., 2017). Collectively, our data show that the class I PI3K pathway is the most potent driver of PI(3,4)P_2_-dependent signaling.

## Results

### cPH probes selectively bind plasma membrane PI(3,4)P_*2*_

The TAPP1 protein (encoded by the *PLEKHA1* gene) contain both N- and C-terminal PH domains (figure 1A), the latter of which selectively binds PI(3,4)P_2_ (Dowler et al., 2000). Previous studies using the isolated TAPP1 C-terminal PH domain (cPH) as a lipid biosensor found no detectable translocation in response to stimuli that induce modest PI(3,4)P_2_ accumulation (Kimber et al., 2002). To improve avidity, tandem dimers of TAPP1 fragments containing the cPH domain have been employed (Oikawa et al., 2008; Posor et al., 2013; He et al., 2017). However, these constructs are based on a fragment including the entire C-terminus (Furutani et al., 2006), which carries binding sites for other proteins in its tail, and have been shown to induce dominant negative effects (Kimber et al., 2002; Hogan et al., 2004; Thalappilly et al., 2008; Li and Marshall, 2015). Therefore, we built tandem dimers and trimers of the isolated cPH domain (residues 169-329 in human TAPP1) as previously defined (Marshall et al., 2002), and included a nuclear export sequence (figure 1A).

**Figure 1:**
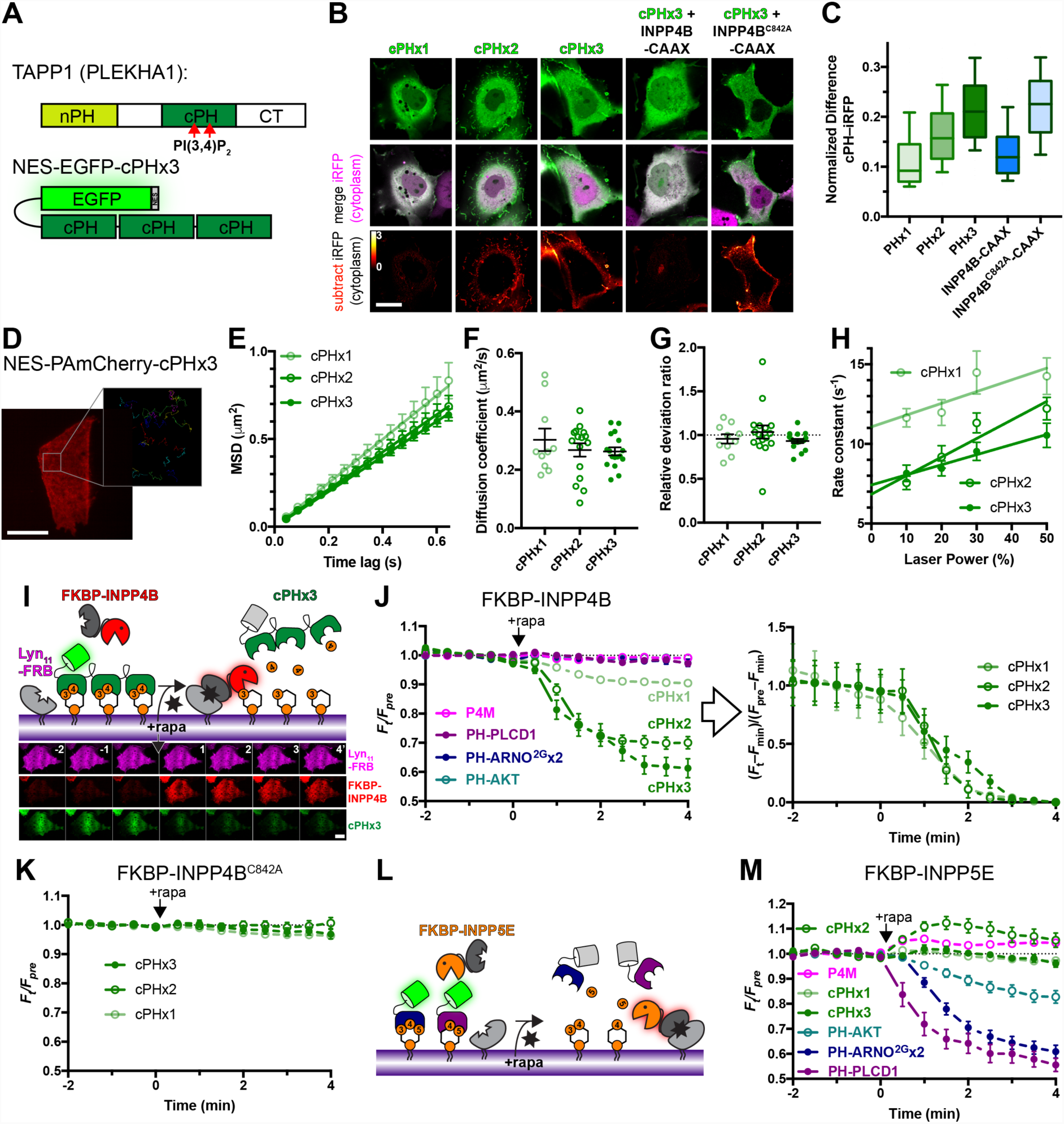
TAPP1 cPHx3 binds PM PI(3,4)P_2_ selectively. (**A**) domain structure of full-length TAPP1 protein, along with the EGFP-tandem cPHx3 fusion incorporating a nuclear export sequence (NES). (**B-C**) **cPHx2 and cPHx3 bind the PM in a PI(3,4)P_2_-dependent manner in COS-7 grown in 10% serum**. Confocal sections (B) are shown of cells expressing EGFP-cPH plasmids and iRFP to mark the cytoplasm. Subtracting normalized iRFP signal from normalized EGFP reveals PM localization of cPHx2 and cPHx3, which is removed by co-expressing CAAX-box tagged PI(3,4)P_2_-phosphatase INPP4B, but not its inactive mutant C842A. Scale bar = 20 µm. The box and whisker plots show median, IQR and 10^th^-90^th^ percentiles of 90-93 cells from three independent experiments. (**D-H**) **Rapid Brownian diffusion of cPH probes**. Image shows COS-7 cell expressing PAmCherry-cPHx3 after photoactivation in TIRF. The inset shows prior trajectories of individual molecules activated at low efficiency, imaged at 16.7 Hz. Scale bar = 20 µm (2 µm for inset). (E) shows mean square displacement vs time lag plots for indicated cPH probes; data are grand means with s.e.m. of 10 (cPHx1), 16 (cPHx2) or 17 (cPHx3) cells. Individual mean cellular diffusion coefficients (F), relative deviation ratio from Brownian motion (G) and “off rate” constant of trajectory lifetimes (from one-phase exponential fit at each laser power, 10-11 cells) are shown with means ± s.e.m. (**I-K**) **Acute PI(3,4)P_2_ depletion removes cPH probes from the PM**. Chemically-induced dimerization (CID) between FKBP and FRB to recruit INPP4B to the PM (I); montage shows TIRF images of a representative COS-7 cell, scale bar = 20 µm. The graphs (J) show normalized fluorescence intensity in TIRF of cells expressing Lyn_11_-FRB-iRFP, mCherry-FKBP-INPP4B and the indicated EGFP-lipid biosensor. Data are means ± s.e.m. of 28-30 cells from three independent experiments. The graph at right is the same data normalized to the maximum change in fluorescence for each cPH construct, to emphasize similar dissociation kinetics. The graph in (K) shows the same experiment for cells expressing a catalytically inactive INPP4B; data are means ± s.e.m. of 27-30 cells from three independent experiments. (**L-M**) **PIP_3_ or PI(4,5)P_2_ depletion does not deplete cPH from the PM**. The same experiment as in (I) is depicted, except 5-phopshatase FKBP-INPP5E replaces INPP4B. The graph in (M) shows mean ± s.e.m. of 28-30 cells from three independent experiments.

Expression of EGFP-tagged cPH monomers, dimers or trimers in COS-7 cells maintained in 10% serum exhibit a predominantly cytosolic localization of the probe when viewed by confocal microscopy (figure 1B). However, comparison with another cytosolic protein (infra-red fluorescent protein, iRFP) revealed enrichment of cPHx2 and cPHx3 at the cell periphery; normalization of the two signals followed by subtracting the iRFP signal from that of EGFP (see Materials and Methods) showed striking cPHx2/3 peripheral localization (figure 1B). This signal was abolished by a PM-targeted INPP4B phosphatase that specifically degrades PI(3,4)P_2_ (Gewinner et al., 2009), though not by a catalytically inactive mutant (figure 1B). Quantification of the difference between EGFP and iRFP signals (figure 1C with Kruskal-Wallis statistic 152.5; *p* < 10^−4^) yielded significantly increasing signal from cPHx1 to x3; INPP4B-CAAX significantly reduced cPHx3 signal (*p* < 10^−4^ in each case, Dunn’s multiple comparison test), whereas the C842A inactive mutant did not (*p* > 0.99). Therefore, cPHx2 and cPHx3 could detect PI(3,4)P_2_ at the plasma membrane of living cells in the presence of serum.

A potential caveat to using tandem arrays of lipid binding domains is that the resulting probe may cluster lipids and exhibit aberrant localization or mobility in the membrane. To determine whether this was the case, we performed single molecule imaging of cPHx1, x2 and x3 mobility in the plasma membrane by tagging with photoactivatable mCherry (PAmCherry). As shown in figure 1D, full activation of the probe with violet light led to uniform labelling of the plasma membrane in total internal reflection fluorescence microscopy (TIRFM), whereas low activation intensities allowed us to image single molecule trajectories (Manley et al., 2008). Analysis of the mean square displacement of these trajectories over time revealed free Brownian motion of all three probes (figure 1E). cPHx1, x2 and x3 diffused with an apparent diffusion coefficient of ∼0.3 µm^2^/s (figure 1F) and did not substantially differ (F = 0.71, *p* = 0.50, one-way ANOVA), and all exhibited a relative deviation ratio close to 1 (figure 1G), meaning there was no substantial difference in actual displacement relative to that predicted by free Brownian diffusion (Fujiwara et al., 2016). Therefore, the tandem array of domains did not slow the probe’s mobility in the membrane.

The lifetime distribution of the single molecules on the membrane followed a single exponential decay with a characteristic “off rate” constant (*k*_off_) that increased linearly with laser power due to increased photobleaching (figure 1H). Extrapolation of this off rate to the intercept at zero illumination power allowed us to estimate the true off rate of the probes. cPHx1 has a lifetime on the membrane of approximately 90 ms (off rate = 11 s^−1^), whereas cPHx2 and cPHx3 both showed lifetimes of approximately 140 ms. This change in dissociation rate of the tandem arrays was comparatively modest, and certainly not multiplicative as would be expected from multi-ligand binding. This implies that, for the majority of probe molecules at steady state, the PH domains are not all occupied by lipid, most likely because of the low abundance of PI(3,4)P_2_. The enhanced PM binding observed in figure 1C likely stems from transient dual or triple occupancy of the PH domains slowing the overall off-rate of the population (and perhaps enhancing the on rate for cPHx3 vs cPHx2).

A potential limitation of biosensors is that when bound to lipid, they will protect the lipid from consumption by enzymes and occlude interaction with endogenous effector proteins, both of which may be needed for local enrichment of the lipid. Therefore, free diffusion of the probe:lipid complex can disrupt local enrichment rather than reporting on it. From the relationship r = (2*D*/*k*off)^0^.^5^ (Teruel and Meyer, 2000), the distance, r, that cPHx3 typically diffuses while attached to lipid can be estimated as ∼290 nm, not much larger than the diffraction limit. This implies there will be a limited propensity of the probes’ free diffusion to “smear out” local enrichment of PI(3,4)P_2_ molecules that are detectable with diffraction-limited optical imaging.

To determine if PM localization of cPH probes is sensitive to acute depletion of PI(3,4)P_2_, we utilized chemically-induced dimerization between FK506 binding protein 12 (FKBP) and the FKBP12 Rapamycin Binding (FRB) domain of mTor (Belshaw et al., 1996) to recruit INPP4B to the PM (figure 1I). Rapamycin-induced dimerization and thus INPP4B recruitment led to depletion of PM-associated cPH signal in TIRFM (figure 1J, two-way ANOVA of area under the curve [AUC] before and after rapamycin addition, *F*(1, 200) = 314.5, *p* < 10^−4^; *p* < 10^−4^ for cPHx1, x2 and x3, Sidak’s multiple comparisons test). The depletion was progressively greater for cPHx3 > cPHx2 > cPHx1 (comparing AUC, *F* = 75.64; *p* < 10^−4^; cPHx2 vs cPHx3, *p* = 0.02; cPHx1 vs cPHx2, *p* < 10^-4^), reflecting the increased avidity of the tandem probes for PM PI(3,4)P_2_. On the other hand, probes for other inositol lipids, including PI4P (P4Mx1), PI(4,5)P_2_ (PH-PLCD1), PIP_3_ (PH-ARNO^2G^x2) or PI(3,4)P_2_/PIP_3_ (PH-AKT) did not show a significant change (Hammond et al., 2014; Várnai and Balla, 1998; Venkateswarlu et al., 1998; Watton and Downward, 1999). Notably, when the data were normalized to emphasize the kinetics of the change, no difference between the rates of cPHx1, x2 or x3 dissociation after INPP4B recruitment were observed (figure 1J, right; Kruskal-Wallis statistic = 1.19, *p* = 0.55), indicating that the higher avidity tandem probes did not effectively sequester lipid from the INPP4B enzyme. Furthermore, recruitment of inactive INPP4B mutant C842A did not lead to substantial depletion of the cPH probes from the PM (figure 1K).

Given the much higher avidity of cPHx3 and x2 for the PM, we wanted to rule out a weak interaction with other lipids that might synergize to influence PM targeting by these probes. To this end, we used chemically induced dimerization again, this time using the INPP5E phosphatase domain (figure 1L), since this domain is active against PI(4,5)P_2_ and PIP_3_, generating PI4P and PI(3,4)P_2_, respectively (Asano et al., 1999; Kong et al., 2000). INPP5E recruitment did not displace PI4P-binding P4Mx1 or any of the cPH probes from the PM, though it did displace probes that bound to PIP_3_ or PI(4,5)P_2_ (figure 1M). Therefore, these two inositol lipids do not influence the PM-association of cPH probes.

Together, these results allowed us to conclude that (i) the cPHx3 probe derived from TAPP1 exhibits significantly greater localization than previously used single and tandem dimer versions, (ii) the tandem trimer configuration does not disrupt free diffusion in the plane of the membrane nor sequester PI(3,4)P_2_ to an extent that prevents INPP4B access and (iii) exhibits localization that depends solely on the phosphoinositide PI(3,4)P_2_. These are crucial criteria in the definition of a reliable genetically encoded lipid biosensor. However, there is another critical criterion that must be met, which we tested next.

### PI(3,4)P_*2*_ is sufficient for cPHx3 cellular localization

We have argued that a crucial but often ignored criterion for a high-fidelity lipid biosensor is a sole requirement of the lipid to drive biosensor localization (Wills et al., 2018). A convenient way to test this criterion is to induce ectopic synthesis of the lipid elsewhere in the cell and test whether this is sufficient to recruit the biosensor (Hammond et al., 2014). To this end, we turned to the *Legionella* effector protein LepB, the N-terminus of which possesses PI3P 4-OH kinase activity, thus generating PI(3,4)P_2_ via a non-canonical pathway not utilized by mammalian cells (Dong et al., 2016). We found that expression of LepB led to substantial PI3P depletion from cells, and consequent swelling of the endosomal compartment that depends on this lipid for function (B.D.G and G.R.V.H., unpublished observations). Therefore, to induce acute LepB activity, we turned to an optogenetic approach that utilizes genetic code expansion to incorporate an unnatural, caged amino acid into the protein (Luo et al., 2014; Liu et al., 2017; Courtney and Deiters, 2018).

This system relies on mutating the target protein at the desired codon to the infrequently used amber stop codon (UAG). The mutant gene is then transfected into cells along with plasmids encoding an engineered pyrrolysyl-tRNA synthetase/tRNA pair and the caged amino acid is added to the media. Here, a hydroxycoumarin-caged lysine (HCK) is transacylated onto the tRNA and thence ribosomally incorporated into the mutated gene in response to the UAG codon. In the context of LepB, the bulky hydroxycoumarin group incorporated onto lysine 39 blocks the active site (Dong et al., 2016). However, illumination of cells expressing this system with 405 nm light causes photolysis of the coumarin group, liberating the lysine residue, and generating wild-type, active LepB (figure 2A). This provides precise and acute spatiotemporal control over LepB function in live human cells.

**Figure 2:**
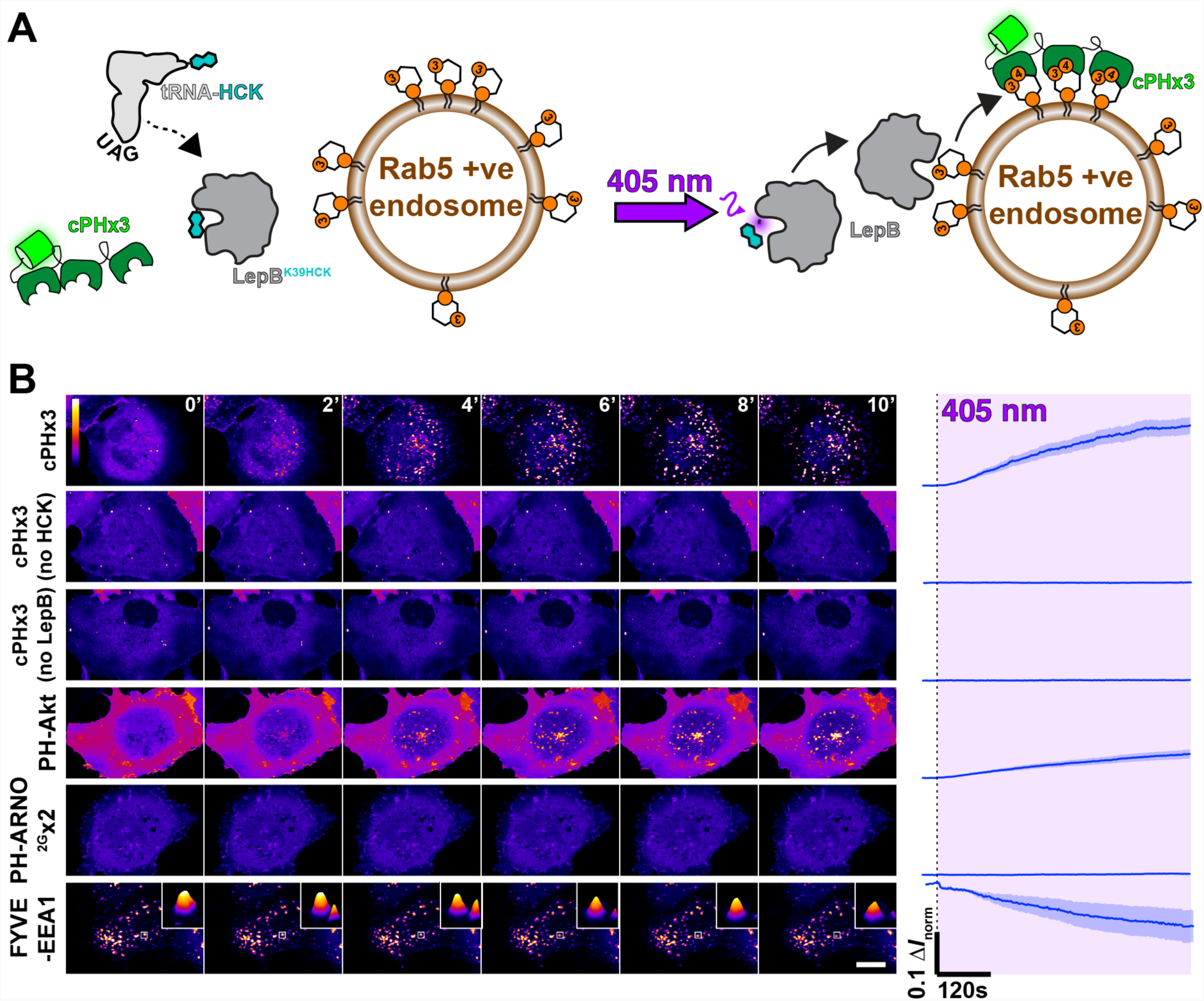
PI(3,4)P_2_ is sufficient to recruit cPHx3. (**A**) **optogenetic activation of PI3P 4-kinase LepB**. Cells are transfected with a plasmid encoding Amber stop codon (UAG) recognizing tRNA and a tRNA synthase incorporating hydroxycumarin lysine (HCK), along with a LepB mutant containing a K39 to UAG mutation, causing incorporation of active-site blocking HCK into the protein. Upon illumination with 405 nm light, the hydroxycumarin is photolyzed, yielding wild-type K39 and inducing PI(3,4)P_2_ synthesis on PI3P-replete endosomes. (**B**) **LepB-synthesis of PI(3,4)P_2_ on endosomes recruits cPHx3 and PH-Akt**, but not the PIP_3_ biosensor PH-ARNO^2G^x2. Images show confocal sections of COS-7 cells transfected with the indicated EGFP-tagged biosensor and the components described in (A) and grown in the presence of HCK. Scale bar = 20 µm. Two controls are shown where either HCK or LepB plasmids were omitted. The graph shows the change in normalized fluorescence intensity in a mask derived from Rab5 puncta (not shown); Trace lines represents mean and shaded area is s.e.m. of 19 (cPHx3), 18 (PH-Akt) or 17 (FYVE and PH-ARNO^2G^x2) cells from four independent experiments or 5 cells (No HCK or no LepB) from a single experiment.

Optogenetic activation of LepB in cells expressing cPHx3 caused synthesis of PI(3,4)P_2_ on endosomes and striking recruitment of the cPHx3 probe (figure 2B). This optogenetic activation was only observed in cells grown in the presence of exogenous HCK, and depended on co-transfection with the LepB^K39UAG^, ruling out off-target effects on endogenous UAG-containing genes or of light-exposure alone. Similarly, we observed clear translocation of PH-AKT to endosomes, which is expected given this PH domain’s binding to both PIP_3_ and PI(3,4)P_2_ (Ebner et al., 2017). However, no recruitment of the PIP_3_-specific PH-ARNO^2G^x2 was observed (Venkateswarlu et al., 1998; Manna et al., 2007), demonstrating that there was no further phosphorylation of the lipid at the 5-OH. Finally, we observed some depletion of the PI3P biosensor FYVE-EEA1 (Wills et al., 2018), though the biosensor remained mostly endosome-bound, indicating a small fraction of total PI3P was converted to PI(3,4)P_2_ over the time course of these experiments (figure 2B).

Collectively, these results demonstrate that generation of PI(3,4)P_2_ in a cellular membrane is indeed sufficient to recruit cPHx3, which given its extensive *in vitro* selectivity (Dowler et al., 2000; Thomas et al., 2001), now passes all the key criteria that define a high-fidelity, genetically encoded biosensor for PI(3,4)P_2_ (Wills et al., 2018). The fidelity of the probe established, we next turned our attention to deploying this tool to answer some central questions about the lipid’s metabolism and function.

### Dominance of Class I PI3K in PI(3,4)P_*2*_ synthesis

The canonical view of PI(3,4)P_2_ synthesis induced by Class I PI3K signaling is that PI3K generates PIP_3_, which is then dephosphorylated at the 5-OH position by inositol polyphosphatase 5-phosphatase (INPP5) family members SHIP1 or SHIP2 (Vanhaesebroeck et al., 2012). Both enzymes are activated by tyrosine phosphorylation cascades initiated by the activated receptor tyrosine kinase. The extent of PI(3,4)P_2_ accumulation varies depending on receptor activation, with insulin or insulin-like growth factor reported to generate only modest amounts of the lipid (Jackson et al., 1992; Guilherme et al., 1996), which have not been detected with a single cPH probe (Kimber et al., 2002).

We tested the capacity of cPHx3 to detect insulin-induced PI(3,4)P_2_ production in HeLa cells (Figure 3A). To unambiguously detect PIP_3_ in the same cells, we employed a tandem dimer of the PH domain from ARNO, specifically selecting the 2G splice variant with high PIP_3_ affinity (Cronin et al., 2004) and introducing the I303E mutation that prevents PH domain interaction with Arl GTPases (Hofmann et al., 2007). We selected this domain since it exhibits clear discrimination for PIP_3_ over PI(3,4)P_2_ (Manna et al., 2007). Insulin stimulation of HeLa cells generated a robust, transient increase in PIP_3_ biosensor at the PM, with a lagging, yet more sustained appearance of the PI(3,4)P_2_ biosensor (figure 3B, C) – a clear recapitulation of the original biochemical measurements of PI3K product generation (Stephens et al., 1991; Jackson et al., 1992; Hawkins et al., 1992), apparently supporting the canonical view.

**Figure 3:**
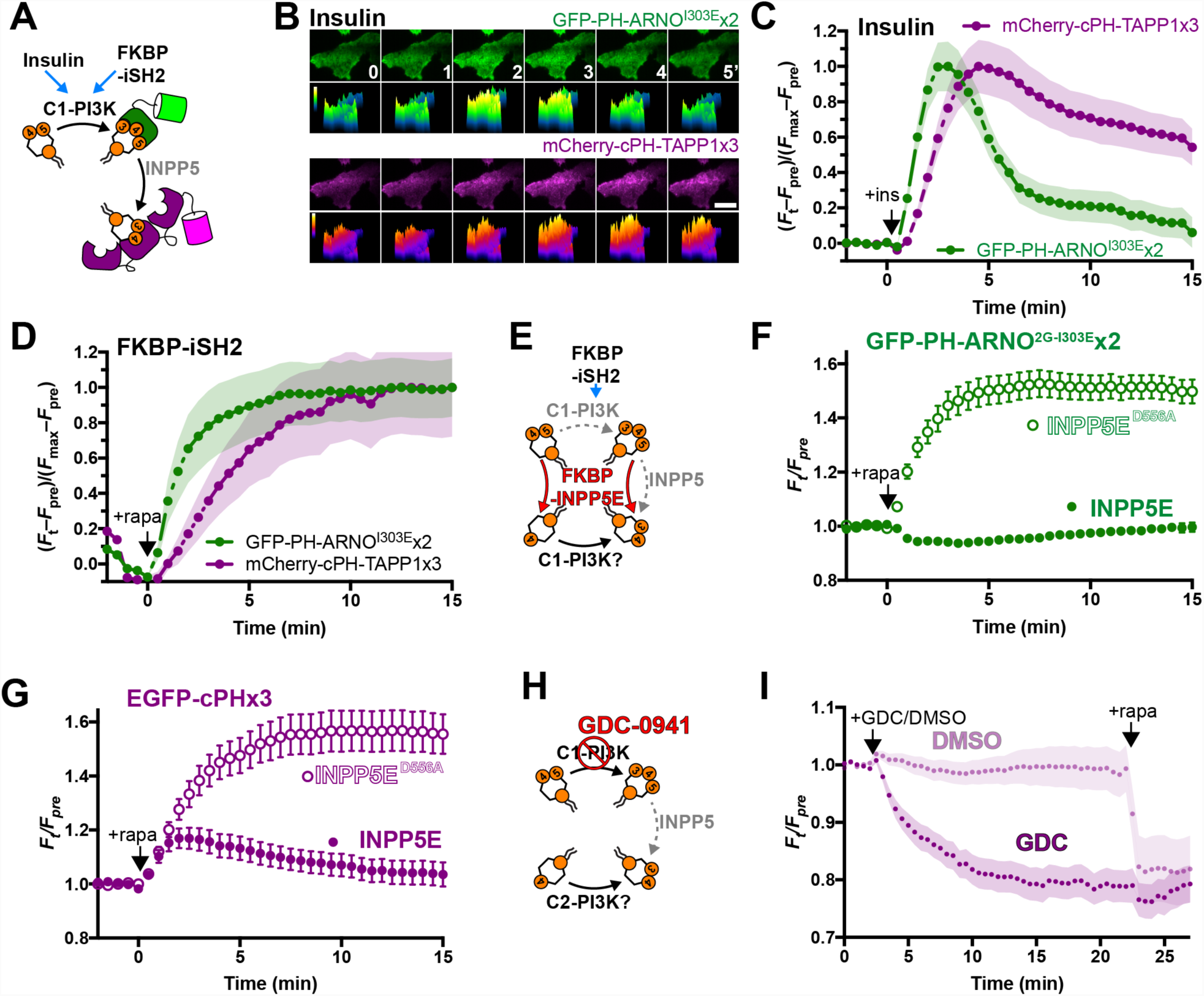
PM PI(3,4)P_2_ is derived from PIP_3_ synthesized via the class I PI3K pathway. (**A**) activation of the C1-PI3K pathway via insulin stimulation or recruitment of endogenous p110 class I PI3K subunit via CID-mediated recruitment of FKBP-iSH2. (**B-C**) **Insulin-stimulated transient synthesis of PIP_3_ and sustained lagging synthesis of PI(3,4)P_2_.** TIRF images of HeLa cells (B) expressing PIP_3_ biosensor EGFP-PH-ARNO^2G-I303E^x2 and PI(3,4)P_2_ biosensor mCherry-cPHx3 stimulated after time 0 with 200 nM insulin. Scale bar = 20 µm. The graph (C) shows fluorescence intensity data normalized to minimum and maximum intensities, and are means with s.e.m. shaded for 34 cells from three independent experiments. (**D**) **Artificial activation of class I PI3K with iSH2 also induced lagging synthesis of PI(3,4)P_2_.** Data are from COS-7 cells expressing the indicated biosensors, along with Lyn_11_-FRB-iRFP and TagBFP2-FKBP-iSH2. They are normalized to minimum and maximum intensities, and are means with s.e.m. shaded for 31 cells from three independent experiments. (**E-G**) **PI(3,4)P_2_ is derived from PIP_3_.** (E) The experimental setup is to activate class I PI3K with FKBP-iSH2 whilst simultaneously recruiting the PIP_3_ and PI(4,5)P_2_-depleting enzyme FKBP-INPP5E, or catalytically inactive D556A mutant as control. Depletion of the PI3K substrate PI(4,5)P_2_ leaves 3-phosphorylation of PI4P as the only route of PI(3,4)P_2_ synthesis. Graphs show data from COS-7 cells imaged by TIRF expressing PIP_3_ biosensor EGFP-PH-ARNO^2G-I303E^x2 (F) or PI(3,4)P_2_ biosensors cPHx3 (G) along with Lyn_11_-FRB-iRFP, TagBFP2-FKBP-iSH2 and mCherry-FKBP-INPP5E wild-type or D556A, as indicated. Data are means ± s.e.m. of 32-40 cells from three independent experiments. (**H-I**) **Most PI(3,4)P_2_ is derived from class I PI3K**. (H) The specific inhibitor GDC-0941 can distinguish class I from class II PI3K activity that could directly produce PI(3,4)P_2_. The graph (I) shows normalized fluorescence intensity EGFP-cPHx3 in cells co-transfected with Lyn_11_-FRB-iRFP and mCherry-FKBP-INPP4B, with CID induced at 22 min to deplete remaining PI(3,4)P_2_. 250 nM GDC-0941 or DMSO was added at 2 min. Data are means with s.e.m. shaded for 34 cells from four independent experiments (GDC) or 18 cells from two independent experiments (DMSO).

As an alternative mechanism to activate Class I PI3K, we turned to chemically induced dimerization to recruit the inter-Src Homology 2 (iSH2) domain from the p85 regulatory subunit of PI3K (figure 3A). This system recruits endogenous PI3K p110 catalytic subunits to the membrane, inducing PIP_3_ synthesis (Suh et al., 2006). However, in this situation, tyrosine kinase-mediated activation of SHIP phosphatases is not expected. Nevertheless, iSH2 recruitment to the PM of COS-7 cells induced rapid PIP_3_ synthesis and the same, lagging PI(3,4)P_2_ accumulation (figure 3D). We therefore speculated that, in this context at least, PI(3,4)P_2_ accumulation might be driven by direct phosphorylation of PI4P by p110, given that the enzyme is known to perform this reaction in a test tube (Carpenter et al., 1990).

To test this speculation, we devised an experiment wherein the INPP5E enzyme would be co-recruited to the PM in conjunction with iSH2. Since INPP5E will deplete both PI(4,5)P_2_ and PIP_3_, accumulation of PIP_3_ should be prevented (figure 3E). PIP_3_ will be degraded into PI(3,4)P_2_ – but synthesis of PI(3,4)P_2_ will only be sustained if the p110 enzymes indeed directly converts PI4P into PI(3,4)P_2_. Compared with controls using a catalytically impaired INPP5E D556A mutant, INPP5E completely blocked the accumulation of the PIP_3_ biosensor at the PM after co-recruitment (figure 3F). However, cPHx3 initially recruited to the PM at a similar rate to control, though its synthesis was rapidly cut off when INPP5E was co-recruited (figure 3G). Therefore, PI(3,4)P_2_ synthesis cannot be sustained in the absence of PIP_3_ generation, even under conditions where SHIP phosphatases are not predicted to become activated. Presumably, the PIP_3_ 5-phosphatase activity responsible either comes from basal SHIP activity through interaction of its C2 domain with acidic PM lipids (Ong et al., 2007; Coq et al., 2017), or else from other INPP5 family members, which are all competent at converting PIP_3_ to PI(3,4)P_2_ (Trésaugues et al., 2014). Identification of the enzymes responsible will be a key question for future work.

Thus far, these experiments addressed the pool of PI(3,4)P_2_ generated downstream of Class I PI3K signaling. However, activity of class II PI3K have been shown to function in a more constitutive capacity (Posor et al., 2013; Marat et al., 2017). We therefore wanted to identify the source of PI(3,4)P_2_ that we observed at the PM in growing cells (figure 1). To this end, we employed the potent and Class I PI3K-selective inhibitor GDC-0941 (Kong et al., 2010) at 250nM to distinguish Class I and Class II activities (figure 3H). After inhibitor treatment for 20 min, cells were subject to chemically-induced dimerization to recruit a co-expressed FKBP-INPP4B to the PM (figure 3I), the goal being to degrade any remaining PI(3,4)P_2_. As shown in figure 3I, GDC treatment led to the depletion of almost the entire pool of PM-associated PI(3,4)P_2_ from cells grown in serum. We cannot rule out that the small additional decrease induced in the 5 min after rapamycin addition is not an addition artifact (figure 3I); area under the curve measurements for post rapamycin treatment vs. the 5 min period preceding it do not show a significant change (*p* = 0.43, Sidak’s multiple comparisons test) as compared to the change following DMSO treatment (*p* < 10^−4^, Sidak’s multiple comparison test after 2-way repeated-measures test, *F* = 61.9, *p* < 10^−4^). However, the data do clearly indicate that the overwhelming majority of PI(3,4)P_2_ present in the PM of cells is generated by the class I PI3K pathway.

### Direct Hydrolysis of PI(3,4)P_*2*_ by PTEN

So far, our data support a canonical view of Class I PI3K signaling, which is dominated by conversion of PI(4,5)P_2_ to PIP_3_, followed by degradation back to PI(4,5)P_2_ by PTEN or conversion to PI(3,4)P_2_ via INPP5 enzymes, most prominently SHIP1 and SHIP2 (Vanhaesebroeck et al., 2012; Fruman et al., 2017). PTEN and INPP5 thus represent a bifurcation of the pathway: in the former case, towards a simple inactivation; in the latter, the conversion to a modified but still active signaling state (Li and Marshall, 2015). PI(3,4)P_2_ is ultimately degraded to PI3P by INPP4A/B, a process intimately linked to endocytosis (Shin et al., 2005). However, it has been perplexing that the accumulation of PI(3,4)P_2_ seen after PI3K activation does not lead to a detectable increase in PI3P levels (Jackson et al., 1992; Stephens et al., 1991), implying an alternative route of degradation. Recently, Malek and colleagues have proposed that PTEN in fact directly converts PI(3,4)P_2_ to PI4P, terminating this signaling as well (Malek et al., 2017). The evidence was based on the requirement for PTEN knockout for EGF-stimulated PI(3,4)P_2_ accumulation, in addition to showing that a PI(3,4)P_2_ 3-phosphatase activity in MCF10 cell cytosol was lost in PTEN nulls. However, direct evidence for hydrolysis of PI(3,4)P_2_ in intact, living cells is still lacking.

The principle reason for the ongoing ambiguity over cellular activity of PTEN against PI(3,4)P_2_ has been the ambiguity when interpreting changes in lipid levels in cells. When PTEN loss induces PI(3,4)P_2_ accumulation, this can be explained by failure of PTEN to degrade PI(3,4)P_2_ directly – or alternatively, due to impaired PIP_3_ degradation, leaving more substrate available for the INPP5 enzymes (figure 4A). Careful mathematical modeling suggested the latter explanation did not explain the accumulation in MCF10 cells (Malek et al., 2017). Nonetheless, the evidence is indirect. We therefore devised an experiment utilizing the LepB PI3P 4-OH kinase. Since we showed that this enzyme generates an endosomal pool of PI(3,4)P_2_ devoid of PIP_3_ (figure 2), activity of PTEN leading to depletion of PI(3,4)P_2_ can be unambiguously assigned to direct hydrolysis of the lipid in this context (figure 4B).

**Figure 4:**
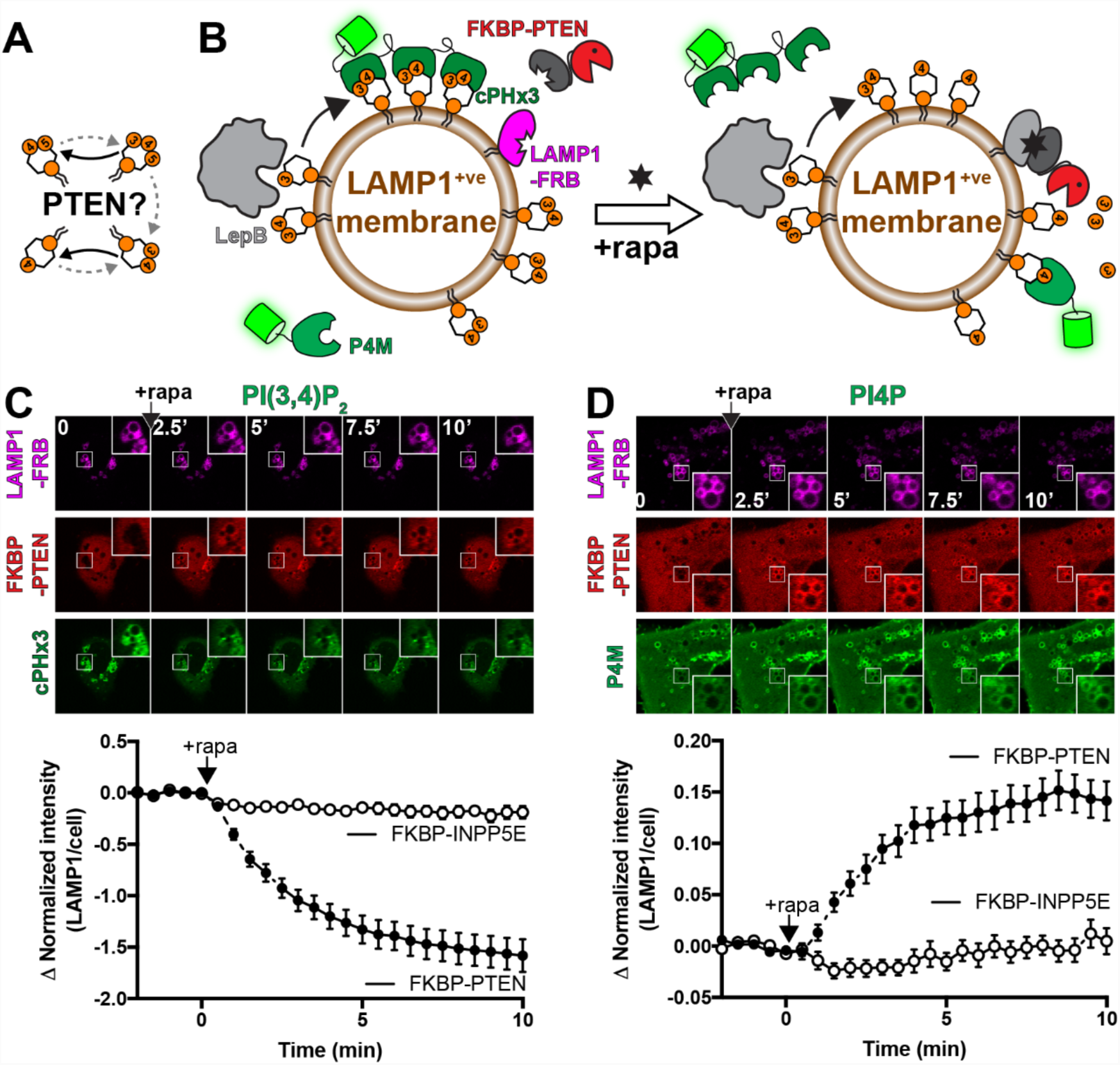
PTEN directly dephosphorylates PI(3,4)P_2_. (**A**) Potential PTEN-driven reactions. (**B**) **Experimental setup**. COS-7 cells are expressing active LepB to drive PI(3,4)P_2_ in the endosomal system, including (but not limited to) LAMP1-positive membranes. LAMP1-FRB is then used for CID-mediated recruitment of mCherry-FKBP-PTEN to the membranes, where changes in lipids are detected with the indicated biosensors. (**C-D**) **PTEN-mediated conversion of PI(3,4)P_2_ to PI4P**. Images show confocal sections of COS-7 cells expressing the indicated constructs before and at the indicated times after CID with rapa. Insets are 5 µm. Data in the graphs are means ± s.e.m. of 32-37 cells from three independent experiments. FKBP-INPP5E serves as a negative control.

Indeed, use of chemically-induced dimerization of FKBP-PTEN to FRB-LAMP1-positive endosomes/lysosomes caused a rapid depletion of endosomal PI(3,4)P_2_ detected with cPHx3 in LepB-expressing COS-7 cells (figure 4C). That this was truly PI(3,4)P_2_ to PI4P conversion induced by PTEN was further evidenced by a clear accumulation of PI4P on these membranes (figure 4D). Therefore, our data provide a direct demonstration of PI(3,4)P_2_ hydrolysis by PTEN in intact, living cells – and confirms the novel role of this enzyme in terminating PI(3,4)P_2_ signaling in addition to signals driven by PIP_3_ (Malek et al., 2017).

## Discussion

In this study, we report a high avidity probe for PI(3,4)P_2_, cPHx3-TAPP1, which satisfies three crucial criteria for a genetically encoded lipid biosensor (Wills et al., 2018): (i) binding of the PH domain is exquisitely specific for PI(3,4)P_2_ (Dowler et al., 2000; Thomas et al., 2001; Manna et al., 2007), (ii) PI(3,4)P_2_ binding is absolutely necessary, and (iii) sufficient for localization in living cells (figures 1 and 3). [text] We use this high fidelity and high sensitivity probe to develop the first direct evidence for the canonical pathway of PI(3,4)P_2_ synthesis by class I PI3K (i.e. by PIP_3_ de-phosphorylation, figure 3), first proposed nearly three decades ago (Stephens et al., 1991; Hawkins et al., 1992). We also provide direct evidence for a much more recent proposition (Malek et al., 2017): that the tumor suppressor PTEN also terminates PI(3,4)P_2_ signals in addition to those mediated by PIP_3_ (figure 4). We also generated the first optogenetically activatable lipid kinase, LepB (figure 2).

How PI(3,4)P_2_ signaling couples to different functions, and their relationship (or not) with the class I PI3K pathway, is still poorly understood (Li and Marshall, 2015; Hawkins and Stephens, 2016). This has largely been due to the difficulty in studying PI(3,4)P_2_ in isolation from the other, more abundant phosphoinositides such as PI(4,5)P_2_ and PIP_3_. However, recent advances in mass spectrometry are allowing sensitive detection of this lipid (Malek et al., 2017; Bui et al., 2018). We anticipate that our tools and cPHx3 probe will complement these approaches and greatly accelerate research in this area.

For now, the clearest demonstration is that most PI(3,4)P_2_ accumulating in cells is derived from INPP5 activity on class I PI3K-synthesized PIP_3_. Without stimulation, resting levels of PI(3,4)P_2_ are extremely low (Stephens et al., 1991; Hawkins et al., 1992), with immunocytochemical estimates suggesting that perhaps 40% of this is generated by the class II PI3K-C2α (Wang et al., 2018). The function of the greatly expanded class I PI3K-driven pool remains an ongoing question. Conversion of PIP_3_ to PI(3,4)P_2_ will terminate signals generated by PIP_3_-selective effectors, maintain signals by more promiscuous effectors (like Akt), and initiate signaling by PI(3,4)P_2_-selective effectors (Hawkins and Stephens, 2016). What is the purpose of this tiered lipid signaling system? A clue comes from perhaps the best characterized PI(3,4)P_2_-selective effector protein: the TAPP proteins from which our probe is derived. Mice harboring point mutations in *Plekha1* and *Plekha2* that disrupt PI(3,4)P_2_-binding of the TAPP1 and TAPP2 cPH domains exhibit augmented PI3K signaling through insulin and B-cell receptors (Gray and Alessi, 2011; Landego et al., 2012), suggesting a major function of the lipid is in negative feedback of class I PI3K signaling. Interestingly, PI(3,4)P_2_ is also implicated in endocytosis (Shin et al., 2005; Posor et al., 2013; Boucrot et al., 2015). Therefore, it could be possible that PI(3,4)P_2_ downregulates PI3K signals by triggering internalization of activated receptors.

Such a model may reconcile conflicting data over the generation of PI(3,4)P_2_ in clathrin mediated endocytosis. The class II PI3K-C2α was shown to generate a pool of PI(3,4)P_2_ essential for constitutive endocytosis (Posor et al., 2013), whereas a co-incidence detector probe showed that the enzyme was dispensable for PI(3,4)P_2_ accumulation at clathrin-coated structures (He et al., 2017). Although seemingly at odds, it is worth noting that careful quantitative modeling of lipid dynamics during formation of the clathrin bud found that just enough PI(3,4)P_2_ is produced to recruit its SNX9 effector protein – meaning that the level of free lipid might be extremely low and invisible to even the most sensitive biosensor (Schöneberg et al., 2017). So why was PI(3,4)P_2_ observed at clathrin structures at all (He et al., 2017)? We propose that the greatly expanded pool of class I PI3K synthesized PI(3,4)P_2_ was being detected. Indeed, the cells in He et al were imaged in 5% serum, and most experiments were conducted in SUM159 cells, which have constitutive class I PI3K activation via the oncogenic 1047A mutation in p110α (Barnabas and Cohen, 2013). Intriguingly, PI(3,4)P_2_ synthesis by class I PI3K activity was also found to be essential for clathrin-independent, fast endophilin mediated endocytosis (Boucrot et al., 2015). An attractive model is therefore that PI3K-C2α produces a minor pool of PI(3,4)P_2_ that supports constitutive clathrin-mediated endocytosis, whereas greatly expanded pools downstream of Class I PI3K are required for accelerated clathrin-mediated and independent endocytosis after activation, whilst also sustaining Akt signaling.

A second major finding is that PI(3,4)P_2_ is directly hydrolyzed by PTEN (Malek et al., 2017). This result is surprising, since meticulous work with purified protein found much poorer PTEN activity against PI(3,4)P_2_ vs. PIP_3_ (McConnachie et al., 2003). The discrepancy likely derives from the very different enzyme-substrate interactions in living cells; indeed, turnover numbers for PTEN in living cells are several orders of magnitude higher than in the test tube (McConnachie et al., 2003; Feng et al., 2014). It therefore seems likely that PTEN regulates PI(3,4)P_2_ signaling in addition to PIP_3_. Intriguingly, the recent observations that PTEN reduced signaling from forming clathrin-coated structures and slows their maturation is entirely consistent with the roles of both class I and class II PI3K proposed in the previous paragraph (Rosselli-Murai et al., 2018). Finally, since PTEN’s activity on PI(3,4)P_2_ is the direct reversal of the PI4P conversion mediated by class II PI3K, and since PTEN has recently been found to localize to endosomal structures (Naguib et al., 2015), PTEN seems poised to regulate PI3K-C2β signaling from late endosomes/lysosomes too (Marat et al., 2017).

In summary, we have developed and validated a sensitive genetically encoded lipid biosensor for PI(3,4)P_2_. We show this lipid accumulates in cells primarily in response to class I PI3K activity, via circuitous dephosphorylation of PIP_3_. We also provide evidence for direct hydrolysis of PI(3,4)P_2_ by PTEN. Collectively, these results provide fresh impetus to dissect the physiological and pathological signaling outcomes driven by PI(3,4)P_2_, and provides a powerful new tool to aid in this endeavor.

## Materials and Methods

### Cell Culture and transfection

COS-7 (ATCC CRL-1651) and HeLa (ATCC CCL-2) cells were cultured in DMEM (low glucose; Life Technologies 10567022) supplemented with 10% heat-inactivated fetal bovine serum (Life Technologies 10438-034), 100 units/ml penicillin, 100 µg/ml streptomycin (Life Technologies 15140122) and 1:1000 chemically-defined lipid supplement (Life Technologies 11905031) at 37°C in a humidified atmosphere with 5% CO_2_. They were passaged twice per week by dissociation in TrpLE (Life Technologies 12604039) and diluting 1 in 5. For transfection, cells were seeded in 35 mm tissue culture dishes with 20 mm Number 1.5 cover glass apertures (CellVis) coated with 5 µg fibronectin (Life Technologies 33016-015). Between 1 and 24 hours post seeding, cells were transfected with 1 µg total plasmid DNA pre-complexed with 3 µg lipofectamine2000 (Life Technologies 11668019) in 200 µl Opti-MEM (Life Technologies 51985091) according to the manufacturer’s instructions. Cells were imaged 18-26 h post-transfection. For unnatural amino acid incorporation, medium was supplemented with 250 µM hydroxycoumarin lysine in parallel with the transfection.

### Plasmids

pPAmCherry-C1 (Addgene plasmid 31929) was a kind gift of Vladislav Verkhusha (Subach et al., 2009). EGFP (*Aequorea victoria* GFP with F64L and S65T mutations with human codon optimization), mCherry, *Rhodopseudomonas palustris* bacteriophytochrome BphP2-derived near-infrared fluorescent protein variant iRFP713 and *Entacmaea quadricolor* GFP-like protein eqFP578 variant pTagBFP2 were used in the Clontech pEGFP-C1 and N1 backbones as described previously (Zewe et al., 2018). TAPP1 cPH and LepB sequences were obtained as synthetic gblocks (IDT). All plasmids were verified by dideoxy sequencing. Constructs generated in this study are freely distributed through Addgene. Plasmids were constructed using standard restriction cloning or NEB HiFi assembly (New England Biolabs E5520S), or else obtained from the sources indicated below:

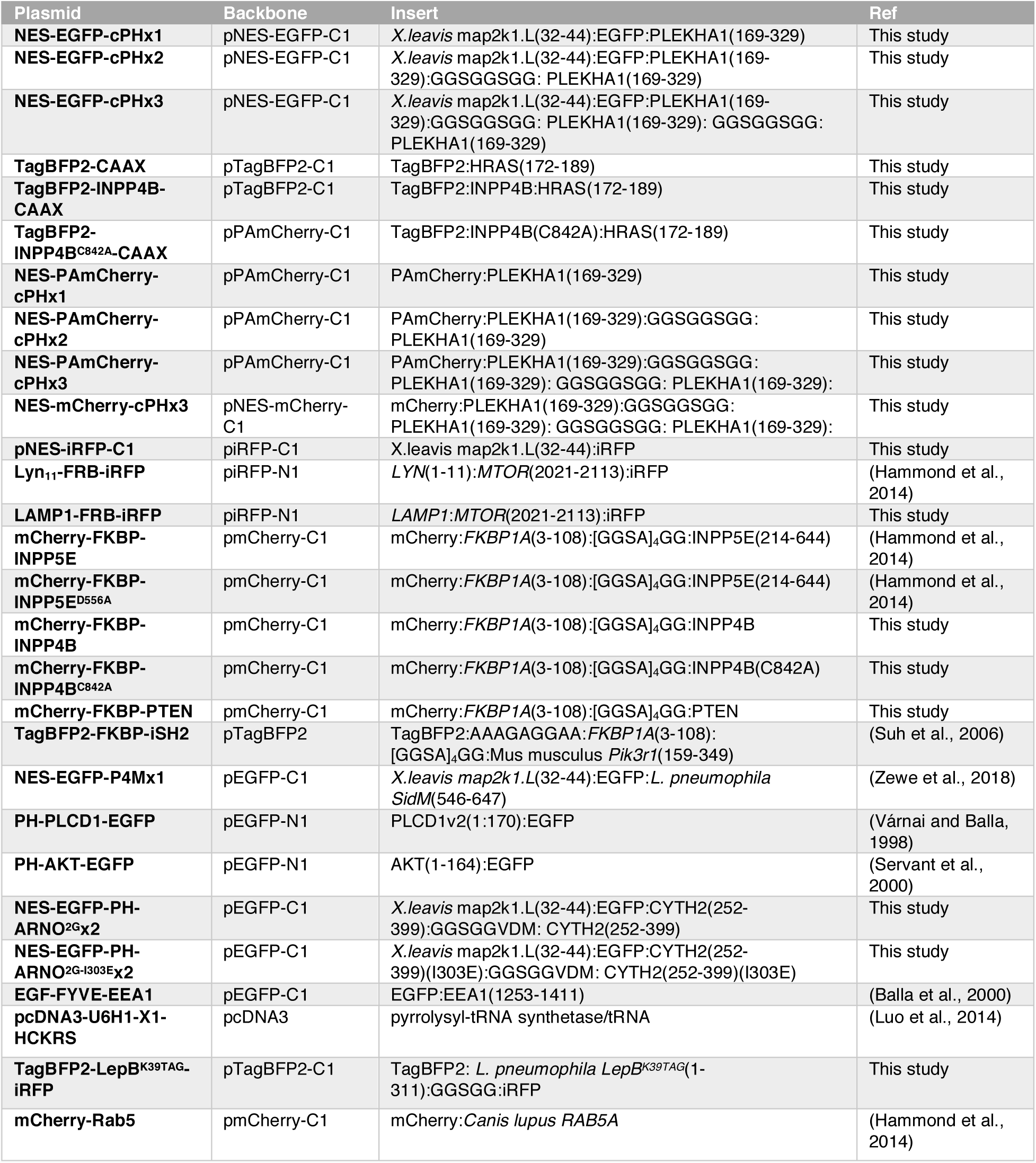

### Chemicals

Rapamycin (Fisher Scientific BP2963-1) was dissolved in DMSO at 1 mM and stored in aliquots at –20°C; the final concentration used in cells was 1 µM. 4 mg/ml Insulin zinc solution (ThermoFisher 12585014) was stored in aliquots at –20°C. GDC-0941 (EMD-Millipore 5.09226.0001) was dissolved in 2 mM DMSO and stored in aliquots at –20°C. Aliquots of hydroxycoumarin Lysine (Luo et al., 2014) were stored at –20°C in DMSO at 100 mM.

### Microscopy

Cells were imaged in 1.6 ml FluoroBrite DMEM (Life Technologies A1896702) supplemented with 25 mM HEPES (pH 7.4) 1:1000 chemically-defined lipid supplement with or without 10% heat-inactivated fetal bovine serum. For treatments, 0.4 ml of this medium containing 5-fold the final concentration of compound was applied to the dish (or 0.5 ml for a second addition).

Confocal microscopy was performed on a Nikon TiE inverted stand with an A1R resonant scan head and fiber-coupled 4-line excitation LU-NV laser combiner equipped with 405, 488, 561 and 640 nm lines. 8 or 16 frame averages were used to improve signal to noise. A 100x 1.45 NA plan-apochromatic oil-immersion objective was used throughout. Blue (405 nm excitation, 425-475 nm emission) and yellow/orange (561 nm Ex, 570-620 nm Em) channels were imaged concurrently, alternating with concurrent imaging of green (488 nm Ex, 500-550 nm Em), far/infra-red (640 nm Ex, 663-737 nm Em) and a transmitted light channel for DIC. The hexagonal confocal pinhole was set to 1.2x Airy disc size of the longest wavelength imaged.

For TIRFM, we used a second Nikon TiE microscope fitted with a TIRF illuminator arm fiber coupled to an Oxxius L4C laser combined equipped with 405, 488, 561 and 640 nm lasers. A 100x 1.45 NA plan-apochromatic oil-immersion objective was used to deliver the high angle of incidence illuminating beam and acquire the images by epi fluorescence. Images were acquired on a Zyla 5.5 sCMOS camera (Andor) with 2x2 binning in rolling shutter mode. Blue (405 nm) and yellow/orange (561 nm excitation) channels were imaged through a dual pass 420-480 & 570-620 nm filter (Chroma), whereas green (488 nm) and far/infra-red (640 nm excitation) used a dual pass 505-550 & 650-850 nm filter (Chroma).

Optogenetic activation of LepB was performed by confocal microscopy. After acquiring approx. 30 s of data with 405 nm illumination set to zero power, transmission was turned up to 20% of the maximum available power from the LU-NV unit.

For single molecule imaging, PAmCherry was imaged without pixel binning in global shutter mode with 25 ms exposures and 30% illumination power with 561 nm, and 0.8% 405 nm for photoactivation from the 100 mW Oxxius lasers. A 16x16 µm region of PM was imaged for tracking.

### Image Analysis

Images in Nikon nd2 image format were imported into the open access image analysis package Fiji (Schindelin et al.), using the LOCI BioFormats importer (Linkert et al., 2010). A custom written macro was used to combine fields into a single enlarged image for the purposes of image analysis (though never for presentation).

### Quantifying the difference in intensities between EGFP and iRFP channels (figure 1B&C)

Regions of interest (ROI) were drawn around each cell, and a custom-written macro was used to measure the average pixel intensity in these ROI and then normalize each pixel to this value. The normalized iRFP (cytosolic) image was then subtracted from the normalized EGFP-cPH channel to yield the subtracted image.

*Intensity changes in specific compartments imaged by confocal (figures 2B, 4B & C):*

Cells were measured inside ROI. A second image channel (mCherry-Rab5 in figure 2 and LAMP1-FRB-iRFP in figure 4) was auto-thresholded and used to generate a mask to measure the normalized pixel intensity of the EGFP channel, as previously described in detail (Zewe et al., 2018).

*PM Intensity changes imaged by TIRFM (figure 1J-M; figure 3):*

ROI corresponding to the footprint of individual cells were defined, and after subtracting camera noise, intensity levels over time were measured. Average pixel intensity in each frame t was normalized to the pre-treatment average level *F*_t_/*F*_pre_.

*Single molecule analysis (figure 1D-H):*

Single molecule trajectories were segmented and tracked using the open-source Fiji implementation of the “u-track” single particle tracking algorithm (Jaqaman et al., 2008). A “Difference of Gaussians” filter was used with a hard threshold and an estimated dimeter of 0.5 µm for the estimated diameter (i.e. an approx 8x8 pixel neighborhood) of single molecules. Co-ordinates of the trajectories were exported as xml files, and mean-square displacement at different time lags (Vrljic et al., 2007) was calculated along with trajectory lifetime distributions using custom written code in Python.

*Selecting representative images for presentation:*

example images were selected based on having the best signal to noise possible, whilst also having measured values close to the sampled population median, and are always within the central interquartile range.

*Data presentation and statistics*

was performed using Graphpad Prism 7. Data were subject to the D’Agostino and Pearson normality test to select for parametric or non-parametric tests.

## Acknowledgements

We thank Vladislav Verkhusha (Albert Einstein College of Medicine) and Tamas Balla (NIH) for generously sharing plasmids. This work was supported by National Institutes of Health grant 1R35GM119412-01 (to G.R.V.H.) and National Science Foundation grant CBET-1603930 (to A.D.)

